# Robust gene coexpression networks using signed distance correlation

**DOI:** 10.1101/2020.06.21.163543

**Authors:** Javier Pardo-Diaz, Lyuba V. Bozhilova, Mariano Beguerisse-Díaz, Philip S. Poole, Charlotte M. Deane, Gesine Reinert

## Abstract

**Motivation:** Even within well studied organisms, many genes lack useful functional annotations. One way to generate such functional information is to infer biological relationships between genes/proteins, using a network of gene coexpression data that includes functional annotations. However, the lack of trustworthy functional annotations can impede the validation of such networks. Hence, there is a need for a principled method to construct gene coexpression networks that capture biological information and are structurally stable even in the absence of functional information.

**Results:** We introduce the concept of signed distance correlation as a measure of dependency between two variables, and apply it to generate gene coexpression networks. Distance correlation offers a more intuitive approach to network construction than commonly used methods such as Pearson correlation. We propose a framework to generate self-consistent networks using signed distance correlation purely from gene expression data, with no additional information. We analyse data from three different organisms to illustrate how networks generated with our method are more stable and capture more biological information compared to networks obtained from Pearson or Spearman correlations.

**Code availability:** https://github.com/javier-pardodiaz/sdcorGCN.

## 1 Introduction

Gene expression data, while noisy, contains key information about biological processes (Kothapalli *et al.*, 2002). Such data are often represented as gene coexpression networks, where nodes are genes and edges represent correlations in their expression across multiple samples (Lee *et al.*, 2004). Representing gene coexpression as networks eases the study and visualisation of the expression data (Weirauch, 2011; Magwene and Kim, 2004). One motivation behind creating these networks is that genes which are coexpressed across multiple samples are likely to have related functions (Hughes *et al.*, 2000; Stuart *et al.*, 2003; van Noort *et al.*, 2003; Makrodimitris *et al.*, 2020), allowing inference of gene function using *guilt by association* approaches (Wolfe *et al.*, 2005). This procedure is especially useful if the studied organism is poorly annotated. For example, *Rhizobium leguminosarum*, a soil bacteria important in agriculture that can infect plants of the legume family and provide them organic nitrogenous compound, has no functional information available for approximately 25% of its predicted genes. Most methods to generate and validate gene coexpression networks use exogenous biological information (such as gene ontologies and metabolic information) to select which edges need to or do not have to be present (e.g.Bar-Joseph *et al.*, 2003; Ihmels *et al.*, 2002; Ucar *et al.*, 2007). Therefore, the lack of reliable genomic functional information may hinder the construction of gene coexpression networks and the validation of their accuracy.

The most widely used methods to generate gene coexpression networks in the absence of exogenous information are based on the absolute value of the Pearson correlation of the expression of gene pairs across samples (Weirauch, 2011). After computing the correlation, there are two alternatives: 1) construct a fully-connected weighted network, or 2) impose a threshold to construct unweighted networks with edges connecting genes whose expression correlation is high enough. The former approach is widely used thanks to the R package WGCNA (Langfelder and Horvath, 2008), but results in noisy networks where gene relationships may not be easy to identify. The latter approach (e.g. George *et al.*, 2019) keeps the strongest relationships; however, it is not obvious which threshold value strikes the right balance. A threshold too low, results in overly dense networks that are difficult to analyse; a threshold that is too high risks discarding valuable information.

A natural way of studying gene expression data is to compare the expression of a gene in different samples to assess how it changes. For any two genes, the most intuitive approach should follow the same straightforward procedure: evaluate how the expression of each gene changes across the different samples and then compare the patterns of changes between the two genes. This same idea underpins the concept of *distance correlation* (Székely *et al.*, 2007). While Pearson correlation measures linear relationships, distance correlation measures the dependence, both linear and nonlinear, between two vectors, and provides a non-negative score that is zero if and only if the vectors are statistically independent (Székely *et al.*, 2007). Thus, distance correlations allow us to identify relationships between the expression of genes beyond linearity; this type of correlation has been successfully used in bioinformatics settings to predict miRNA-disease associations (Zhao *et al.*, 2018) and to generate gene regulatory networks from expression data (Guo *et al.*, 2014; Ghanbari *et al.*, 2018). Gene regulatory networks are different from gene coexpression networks because they are directed, much sparser, and aim to identify regulatory pathways rather than functional associations. We give a detailed definition of distance correlation in the Material and Methods section below.

Here we propose a method to construct gene coexpression networks using distance correlation as an intuitive alternative to networks from Pearson or Spearman correlations. To highlight the strengths of our approach, we construct and compare networks using the three correlations. This work is to our knowledge the first work using distance correlation to construct gene coexpression networks.

Distance correlation is always non-negative; that is, it does not capture whether an association between the expression of two genes is positive or negative. Naturally, this information may be biologically relevant; to overcome this shortcoming we introduce a *signed* distance correlation. After calculating the distance correlation between each pair of genes, we impose a sign, which corresponds to the sign of the Pearson correlation between the expression of the genes.

We construct networks by including only edges between genes for which the signed distance correlation of their expression exceeds a threshold. We select the threshold based on the internal consistency of the networks using the R package COGENT (Bozhilova *et al.*, 2020) instead of using exogenous biological information known (or imposed) *a priori*. We evaluate our method in data from three different organisms. First, we generate an unweighted gene coexpression network for the bacteria *Rhizobium leguminosarum* from microarray data. We then analyse RNA-Seq data from the yeast *Saccharomyces cerevisiae*, and single cell RNA-Seq data from human liver cells. The results of our analysis of yeast and human data can be found in the Supplementary Information.

We evaluate the biological information in our networks using the STRING database (Szklarczyk *et al.*, 2019), which is a protein-protein interaction database with scores for pairs of proteins. The higher the STRING score, the more likely the two proteins interact functionally, physically, or both. Using STRING, we show that networks from signed distance correlation capture more biological information and are structurally more stable than networks based on Pearson or Spearman correlation.

While we apply our method to gene expression data, our method to construct networks from signed distance correlations (in combination with COGENT) can be used in applications beyond bioinformatics.

Data and source code are available from https://github.com/javier-pardodiaz/sdcorGCN.

## 2 Materials and Methods

The method we propose generates an unweighted coexpression network from gene expression data that may come from different sources such as microarrays, RNA-Seq, and single-cell RNA-Seq assays. Fig. 1 illustrates the main steps of the method, which includes data pre-processing, computing correlations, and thresholding to create networks.

**Figure 1:**
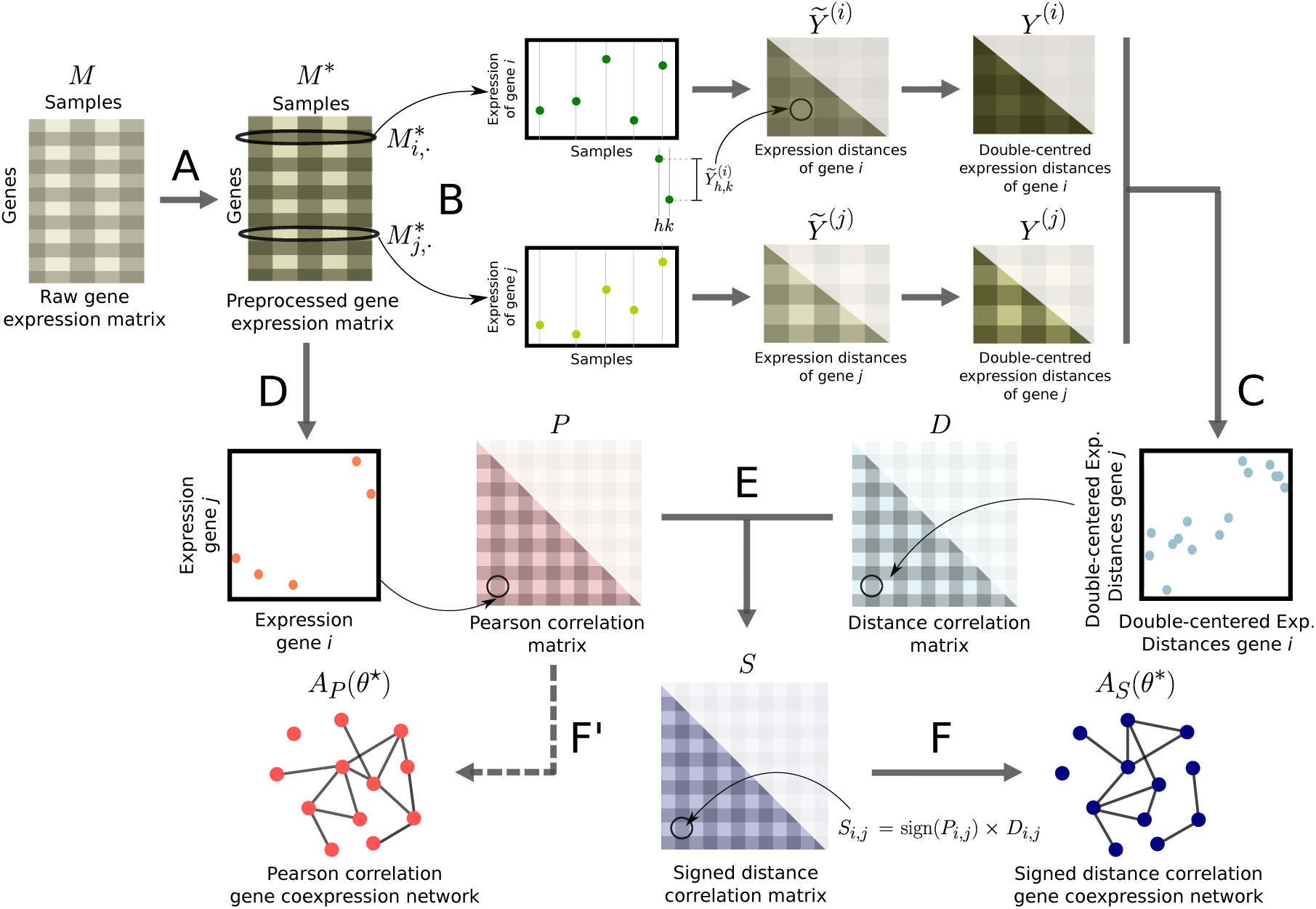
Pipeline to construct networks from gene expression data using signed distance correlation. **A**: We pre-process the input matrix *M* with raw gene expression data using quantile normalisation and setting the lowest 20% values from each sample to the minimum value in *M* to obtain *M* ^*\**^. **B**: We compute the expression distance matrices 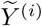 and 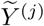 for each gene *i, j* ∈ {1, *… m*,}, and we double center them to obtain *Y* ^(*i*)^ and *Y* ^(*j*)^. **C**: We compute the distance correlation matrix *D*, whose entries *D*_*i,j*_ are the positive root of the Pearson correlation between *Y* ^(*i*)^ and *Y* ^(*j*)^, for every pair of genes. **D**: We compute the Pearson correlation between each pair of rows in the *M* ^*\**^ to obtain the Pearson correlation matrix *P*. **E**: To construct the signed distance correlation matrix *S* we multiply every distance correlation between the expression of two genes *D*_*i,j*_ by the sign of their Pearson correlation sign(*P*_*i,j*_). **F**: Using COGENT (Bozhilova *et al.*, 2020) we find the optimal threshold *θ*^*\**^ that produces the most self-consistent network *A*_*S*_ (*θ*^*\**^) from *S*. **F’**: Analogously, we find the optimal threshold *θ*^⋆^ to generate the network *A*_*P*_ (*θ*^⋆^) from *P*; this step is not part of the pipeline and only necessary to be able to compare Pearson and signed distance correlation networks.

The input to the method is a *m*× *n* gene expression matrix *M*, where each of the *m* rows correspond to a gene, each of the *n* columns is a different sample, and the entries are the expression values of each gene in each sample.

### 2.1 Data

We analyse gene expression data from three different organisms: *R. leguminosarum*, Yeast (*Saccharomyces cerevisiae*), and Human (*Homo sapiens*), obtained using different experimental techniques (microarrays, RNA-Seq and single-cell RNA-Seq). Below we present our results on the *R. leguminosarum* dataset. The description and analysis of yeast and human datasets can be found in Secs. 3 and 4 in the Supplementary Information.

The *R.leguminosarum* bv. *viciae* 3841 data contains gene expression information observed under 18 different growth conditions. These data come from *n* = 54 microarray channels with 3 independent samples per condition (Ramachandran *et al.*, 2011; Pini *et al.*, 2017; Karunakaran *et al.*, 2009). The complete list of the conditions is in Sec. 1 in the Supplementary Information. From the total 7,263 genes in the current genome annotation (Young *et al.*, 2006), we remove genes that do not appear in all the microarrays or appear as pseudogenes, leaving *m* = 7, 077 genes for which we calculate the mean expression within each microarray. These data are encoded in the 7, 077 × 54 matrix *M*.

The data for all three organisms in the form of expression matrices (genes in rows and samples in columns), are available at https://github.com/javier-pardodiaz/sdcorGCN and http://opig.stats.ox.ac.uk/resources.

### 2.2 Pre-processing

Gene expression data is noisy and the raw values are only comparable within the same experiment due to their arbitrary scales. Therefore, gene expression data requires some pre-processing before we can use it to generate networks (Libralon *et al.*, 2009). We apply quantile normalisation (Bolstad *et al.*, 2003) to the gene expression matrix *M*. This normalisation step renders the distribution of the expression values in different samples (i.e., the columns of *M*) identical in their statistical properties, such as maximum value and quantiles. This normalisation enables us to compare data from different experiments.

To avoid interference from low expression values in the quantile normalisation, we ignore the 20% least expressed genes from each sample before the normalisation step. After the quantile normalisation, we set the ignored values to the lowest expression value in *M* to decrease the level of noise. In practice we have observed that 20% offers a good balance between preserving as much information as possible, and weeding out noisy measurements. We denote the pre-processed expression data by *M* ^*\**^, and its *i*-th row by 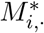.

### 2.3 Computing correlations between the expression of the genes

Distance correlation is as a measure of association between random vectors that addresses some of the limitations of linear measures such as Pearson’s (Székely *et al.*, 2007). To compute the distance correlation between the expression of two genes *i* and *j*, let the vectors 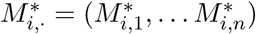 and 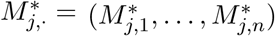 contain their expression values across the *n* samples. For each gene *i* ∈ {1, *…, m*}, we calculate the *n* × *n* expression distance matrix 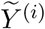 whose entries are the absolute differences between the expression values across samples:

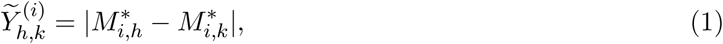

where *h, k* ∈, {1*…, n*} are all the samples in the data. Then, we compute the double-centred expression distance matrix *Y* ^(*i*)^:

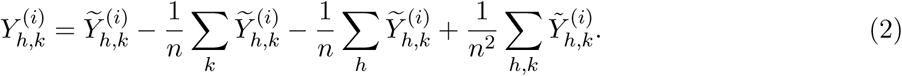

The distance covariance of the expression of genes *i* and *j* is:

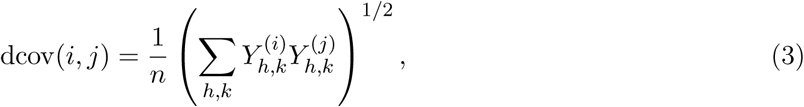

where *h, k* ∈ {1, *…, n*}. Finally, the distance correlation between the expression of the genes is

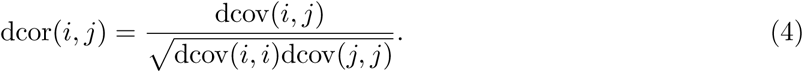

This expression is the non-negative square root of the Pearson correlation between *Y* ^(*i*)^ and *Y* ^(*j*)^. We store the pairwise distance correlation between the expression of all genes in the *m* × *m* symmetric matrix *D* with entries *D*_*i,j*_ = dcor(*i, j*).

The distance correlation in Eq. 4 is always non-negative; however, the association between the expression of two genes can be either positive (i.e., both genes are expressed at the same time) or negative (i.e., one gene is expressed when the other is not). Thus, the sign of the association may contain crucial biological information that is lost if we only use distance correlation. We introduce sign into distance correlation by using the sign of the Pearson correlation matrix *P* between the expression of the genes (which also has size *m* × *m*). Unlike *D*, the correlations in *P* may have negative values; we generate a signed distance correlation matrix *S* whose values are the entries in *D* multiplied by the corresponding entries in *P*:

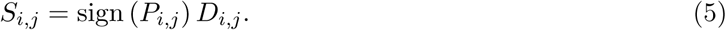

### 2.4 Thresholding correlation matrices

Once we have a signed distance correlation *S*, we generate an unweighted network with adjacency matrix *A*_*S*_(*θ*) by applying a threshold *θ* to *S*:

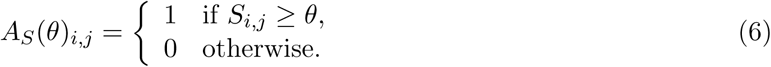

The matrix *A*_*S*_(*θ*) encodes an unweighted undirected network in which pairs of genes are connected if there is a strong and positive signed distance correlation in their expression. Naturally, different values of *θ* result in networks with different properties. We want to find the *θ* that minimises the number of edges between genes that are not coexpressed, and maximises the number of edges between genes that are coexpressed. For this purpose, we use the R package COGENT (Consistency of Gene Expression NeTworks) (Bozhilova *et al.*, 2020).

The main COGENT functions evaluate the internal consistency of a method to generate networks from a specific dataset. First, COGENT splits the gene expression data in two possibly overlapping groups of samples, and then constructs a network with the same node set from each group, *G*_1_ and *G*_2_. Then, COGENT measures the similarity between *G*_1_ and *G*_2_; the more similar the networks, the higher the internal consistency of the method. COGENT helps to find a threshold *θ*^*\**^ that results in the most internally-consistent networks. The COGENT function `getEdgeSimilarityCorrected` provides a score of edge similarity between *G*_1_ and *G*_2_ and adjusts it so that results obtained for different values of *θ* are comparable. We use the semi-random density adjustment implemented in COGENT and select the function parameter that allows to keep the isolated nodes during the analysis.

The edge similarity between *G*_1_ and *G*_2_ is the Jaccard index of the set of edges. This index is adjusted using the randomised networks 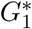 and 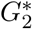 from a a configuration-type model from the degree sequences 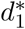 and 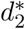. The degree sequences 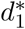 and 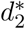 are random permutations of the degree sequences of *G*_1_ and *G*_2_. More precisely, the similarity between *G*_1_ and *G*_2_ is:

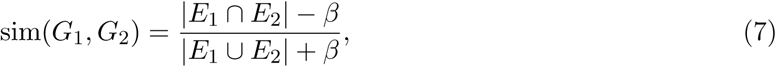

where

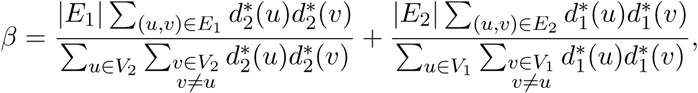

and *E*_*i*_, *V*_*i*_ are the set of edges and vertices for network *G*_*i*_, *i* = 1, 2.

The value of *β* is the expected edge overlap between *G*_1_ and *G*_2_ if they were random networks. In general, *β* is higher for denser networks. The similarity sim(*G*_1_, *G*_2_) in Eq. 7 is a value between −1/3 and 1; this value is high if *G*_1_ is more similar to *G*_2_ than to a randomization of *G*_2_ and vice versa (Bozhilova *et al.*, 2020).

In our analysis, we create two random subsets of columns from *M* to obtain two 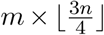 matrices *M*_1_ and *M*_2_, such that half of the total number of samples *n* (i.e., the columns of *M*) are shared between both subsets, and 1/3 of their columns are different. Here 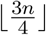 denotes the greatest integer less or equal to 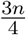. We choose this amount of overlap to evaluate how the relatively small change of about 1/3 of the data affects the final result. Our *in silico* experiments show that if the proportion of samples from *M* shared between *M*_1_ and *M*_2_ is much smaller than 50%, the similarity between the obtained networks is low, and proportions much larger than 50% produce almost identical networks. We pre-process *M*_1_ and *M*_2_ and compute the signed distance correlation matrices *S*_1_ and *S*_2_ as outlined in Sec. 2.3. We test different values of *θ*, and for each of them we obtain two unweighted networks and their similarity with COGENT. We repeat this whole process 25 times, every time with different subsets of columns of *M*. Finally, we compute the similarity score *s*(*θ*), which is the average of the similarity of the networks (in Eq. 7) over the 25 samples.

To favour signal over noise we create a score that balances the similarity of the networks in *s*(*θ*) with the density of *A*_*S*_(*θ*):

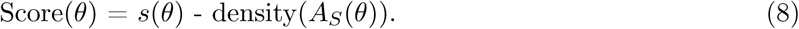

We select the threshold *θ*^*\**^ that retrieves the highest Score(*θ*) and use this value to generate the unweighted gene coexpression network *A*_*S*_(*θ*^*\**^) from the signed distance correlation matrix *S*. We also calculate in the same manner the optimal threshold *θ*^⋆^ to construct an unweighted network *A*_*P*_ (*θ*^⋆^) from the Pearson correlation matrix *P*.

### 2.5 Network comparison

We compare gene coexpression networks obtained from thesholding the signed distance correlation matrix *S* and the Pearson correlation matrix *P*. Letting the edge density of *A*_*S*_(*θ*^*\**^) be *d*_*S*_, and the edge density of *A*_*P*_ (*θ*^⋆^) be *d*_*P*_, we construct two extra networks to obtain a network from each correlation matrix with each edge density. In other words, we compare the following four networks:

- *NS*(*d*_*S*_): Network from *S* with edge density *d*_*S*_ (i.e., *A*_*S*_(*θ*^*\**^)).
- *NP* (*d*_*P*_): Network from *P* with edge density *d*_*P*_ (i.e., *A*_*P*_ (*θ*^⋆^)).
- *NS*(*d*_*P*_): Network from *S* with edge density *d*_*P*_.
- *NP* (*d*_*S*_): Network from *P* with edge density *d*_*S*_.

To construct *NS*(*d*_*P*_) and *NP* (*d*_*S*_), we simply find a threshold manually that produces networks with density *d*_*P*_ and *d*_*S*_.

We first evaluate the internal consistency of each network with COGENT. We also evaluate the biological information contained in the networks using STRING, a database of known and predicted protein–protein interactions (Szklarczyk *et al.*, 2019). STRING collects information from numerous sources, including experimental data, computational predictions and textmining. The association evidence in STRING is categorized into independent channels, weighted, and integrated to produce a confidence score *C* for all recorded protein interactions. Interactions with high *C* score are more likely to be true than those with a low score.

We work with three different sets of confidence scores:

- *C*: Total scores provided by STRING.
- *C*^†^: Scores that *only* consider coexpression information (coexpression channel combined with coexpression transferred channel).
- *C*^‡^: Scores that *exclude* coexpression information.

We provide details of how to retrieve *C*^†^ and *C*^‡^ in the Sec. 5 in the Supplementary Information. For each network we add the confidence score associated with each pair of connected genes. We perform this operation independently for the three sets of confidence scores to obtain three aggregate confidence scores per network. These scores represent the amount of biological information that the networks capture. Then we compare these scores to the expected amount of biological information captured by chance. To do so, we generate two set of 30 random networks with edge densities *d*_*S*_ and *d*_*P*_ and evaluate their biological content following the approach detailed above. We compare the distribution of the aggregate *C, C*^†^ and *C*^‡^ scores from the random networks with density *d*_*S*_ with the scores from networks *NS*(*d*_*S*_) and *NP* (*d*_*S*_) and the scores from the random networks with density *d*_*P*_ with *NS*(*d*_*P*_) and *NP* (*d*_*P*_). In the Sec. 2 in the Supplementary Information, we also perform the biological evaluation of the networks *NR*(*d*_*S*_) and *NR*(*d*_*P*_) with edge density *d*_*S*_ and *d*_*P*_ respectively obtained from a Spearman correlation matrix *R*.

## 3 Results

Here we present our analysis of the *R. leguminosarum* dataset; the results for the yeast and human data are in Secs. 3 and 4 in the Supplementary Information.

### 3.1 Correlation Matrices

The correlation matrices *P* (Pearson), *D* (distance), and *S* (signed distance) are all symmetric with *m* = 7, 077 rows and columns. As we show in Table 1 and Fig. 2, the distribution of the absolute values in *P* are different to those in *D*. In particular, the distribution of the values of *P* and *S* are different. Figs. S2 and S5, and Tables S4 and S7 in the Supplementary Information contain the same analysis for the yeast and human data.

**Table 1:**
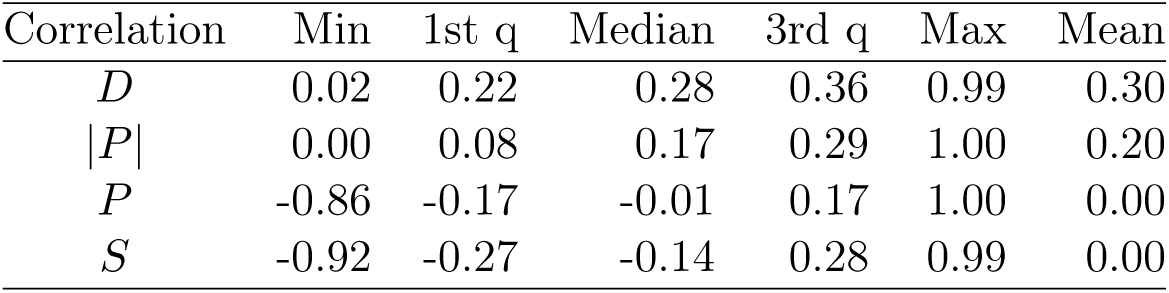
Statistical summary of the correlation matrices from the *R. leguminosarum* data.

**Figure 2:**
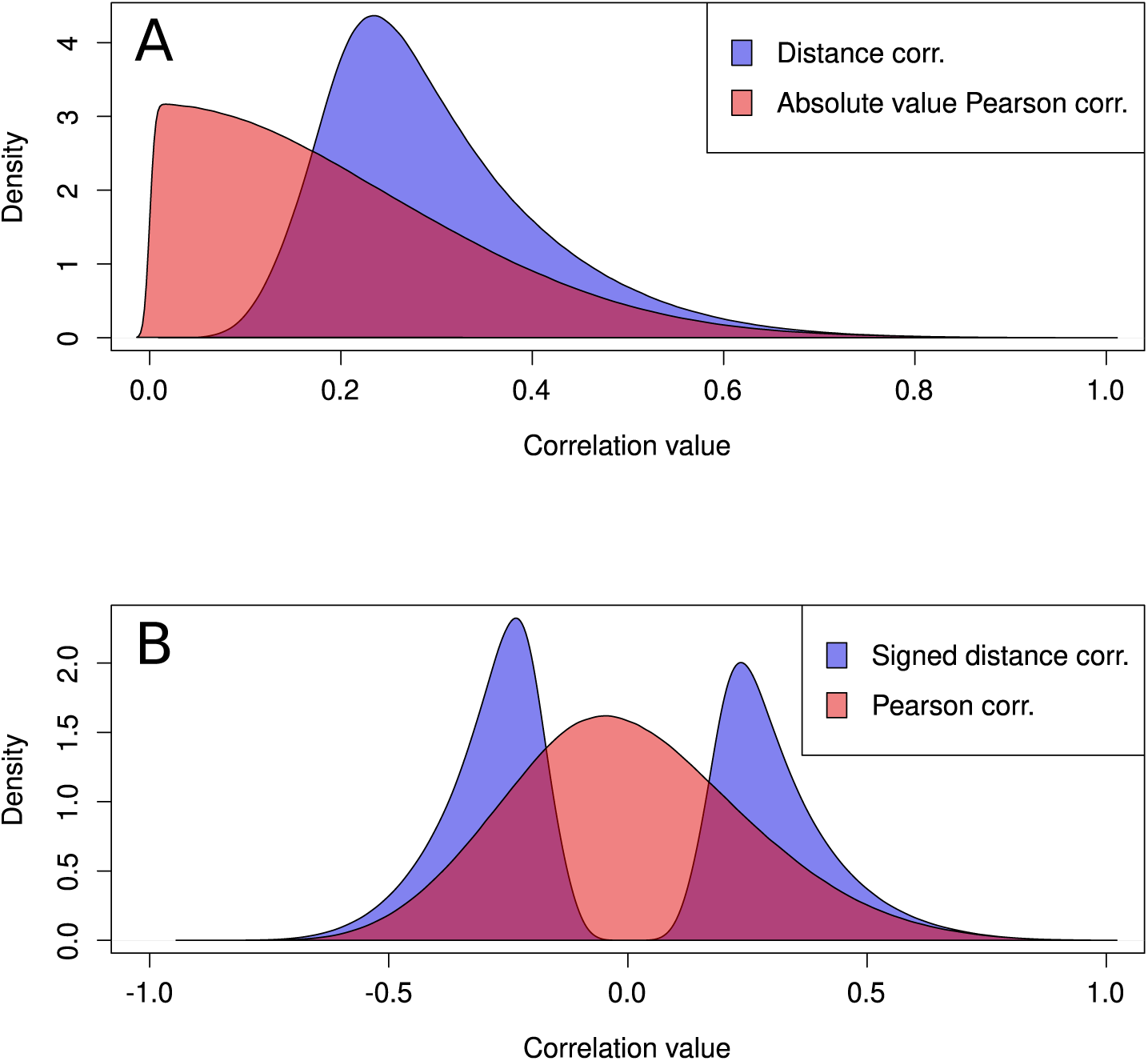
Distribution of the entries of the correlation matrices from the *R. leguminosarum dataset*. **A**: Distribution of distance correlations in *D* (blue), and the absolute of the Pearson correlations *|P|* (red). **B**: Distribution of signed distance correlation *S* (blue), and Pearson correlation *P* (red).

### 3.2 Gene Coexpression Networks

We first estimate, using COGENT (Bozhilova *et al.*, 2020), the optimal thresholds *θ*^*\**^ and *θ*^⋆^ to construct the networks *NS*(*d*_*S*_) and *NS*(*d*_*P*_) from the correlation matrices *S* and *P*, respectively. To analyse and compare the networks, it is perhaps more intuitive to compare the networks using their edge density than with the threshold used to produce them. For this, we construct networks for a range of values of *θ*. For each density (and the *θ* that produced it), we evaluate its consistency score Score(*θ*) using Eq. 8. This score is related to the self-consistency of the network. Fig. 3 shows the value of Score(*θ*) as a function of the density of the networks.

**Figure 3:**
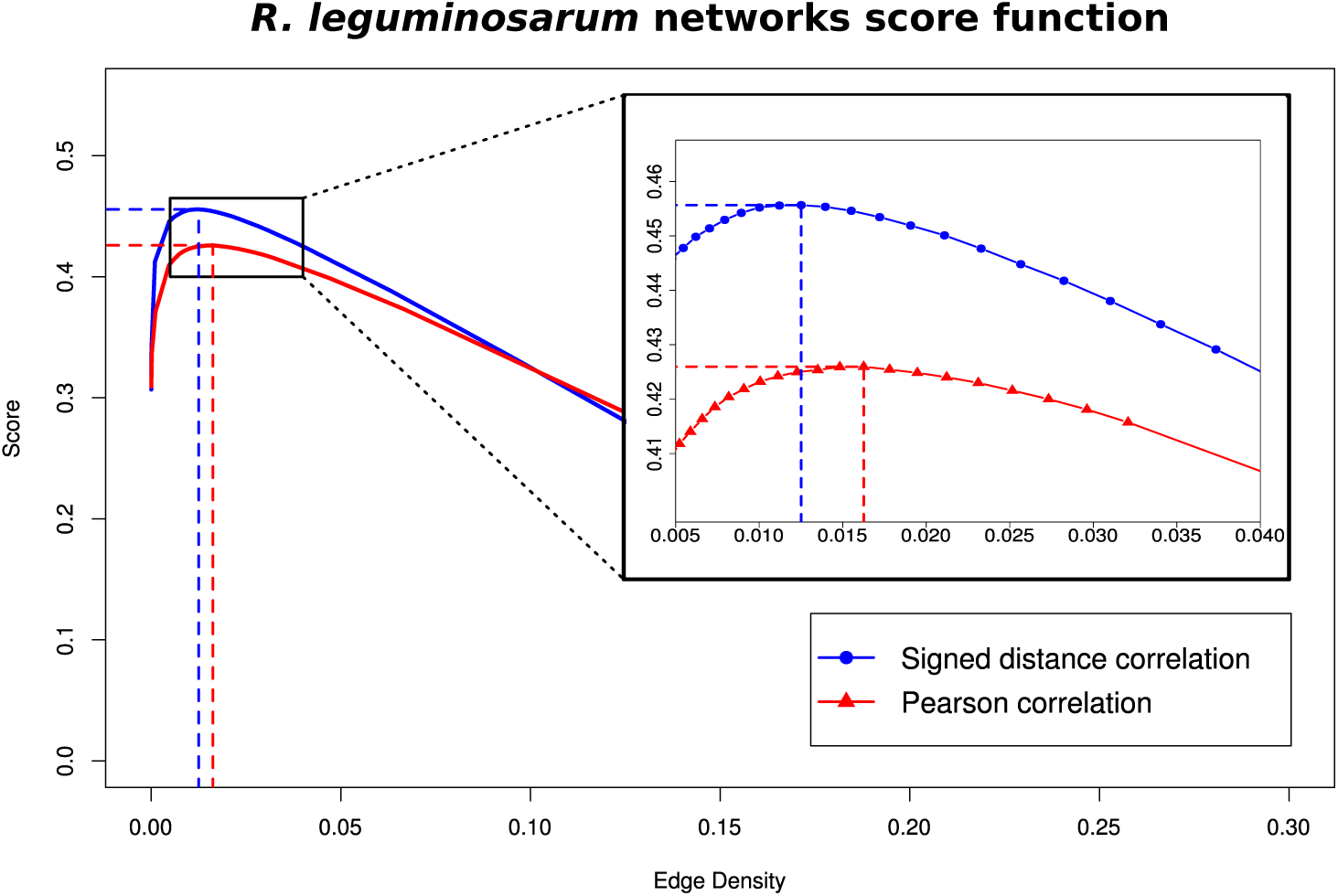
Self-consistency scores of *R. leguminosarum* networks with edge densities between 0 and 0.3. The blue line with circles shows the scores of networks obtained using signed distance correlations; the red line with triangles shows the score of networks using Pearson correlations. The dashed vertical lines indicate the density of the most self-consistent network for each type of correlation.

The highest score from the signed distance correlation networks *NS*(*d*_*S*_) is Score(*θ*^*\**^) = 0.456, where *θ*^*\**^ = 0.62, and density *d*_*S*_ = 1.25%. The highest score from the Pearson correlation networks *NP* (*d*_*P*_) is Score(*θ*^⋆^) = 0.426, where *θ*^⋆^ = 0.58, and density *d*_⋆_ = 1.63%. We also create a network *NP* (*d*_*S*_) from *P* to match the edge density of *NS*(*d*_*S*_), and *NS*(*d*_*P*_) from *S* to match the edge density of *NP* (*d*_*P*_). Table 2 contains a statistical summary of the networks. For both edge densities, the networks retrieved using our signed distance correlation matrix *S* (*NS*(*d*_*S*_) and *NS*(*d*_*P*_)) have a smaller and denser largest connected component (LCC) than the networks obtained using Pearson correlation (*NP* (*d*_*S*_) and *NP* (*d*_*P*_)). As a measure of global clustering we include the global clustering coefficient. See Tables S5 and S8 in the Supplementary Information for the gene coexpression networks for the yeast and human data.

**Table 2:** Statistical summary of *R. leguminosarum* coexpression networks. LCC denotes largest connected component.

| Network | Number of Edges | Number of vertices in LCC | Edge density LCC * 100 | Global clustering coeff. LCC |
| --- | --- | --- | --- | --- |
| $NS(d_S)$ | 313,348 | 6,431 | 1.52 | 0.570 |
| $NS(d_P)$ | 406,977 | 6,688 | 1.82 | 0.570 |
| $NP(d_S)$ | 313,348 | 6,697 | 1.40 | 0.544 |
| $NP(d_P)$ | 406,977 | 6,880 | 1.72 | 0.543 |

### 3.3 Network evaluation

We perform two evaluations of the networks from signed distance correlation and Pearson correlation: self-consistency and biological content. We are interested in the self-consistency of the networks because it reflects their ability to cope with changes in the data used to generate them. If our network is more self-consistent, then we can have greater confidence in the biological conclusions we draw from it, even if the data is imperfect or noisy. The Jaccard similarity in Eq. 7 measures the similarity between networks generated from overlapping, non-identical subsets of a dataset using the same network construction method. The higher the similarity, the higher the internal consistency of the method and the more self-consistent the network obtained from applying it is. We measure the self-consistency Score(*θ*), obtained from computing the average Jaccard similarity of networks from different randomised subsets of data and subtracting the density of *A*_*S*_(*θ*) (*A*_*P*_ (*θ*) for Pearson correlation). As we show in Fig. 3, the self-consistency scores of the networks from signed distance correlations are consistently higher than in the Pearson networks over a an interval of densities that includes the maxima for both correlations. From this analysis we conclude that signed distance correlation networks are more self-consistent than networks based on Pearson correlation. Even the optimal threshold for the Pearson matrix *P* produces a less self-consistent network *NP* (*d*_*P*_) than a signed distance correlation network with a matching edge density (*NS*(*d*_*P*_)). See Figs. S3 and S6 in the Supplementary Information for the same analysis on the yeast and human data, with similar results.

To evaluate the biological content of a network, we add the STRING scores of the edges using: all the information in STRING (*C*), only coexpression information (*C*^†^), and everything except coexpression information (*C*^‡^), as described in Sec. 2.5. Table 3 and Fig. 4 (and Fig. S1 in the Supplementary Information) show the results for the four networks, and the mean scores from random networks. In every case, the signed distance correlation networks contain more biological information than networks from Pearson correlations and the randomised networks. See Tables S6 and S9 and Fig. S4 and Fig. S7 in the Supplementary Information for a similar analysis on the human and yeast data, with a similar results. The highest difference of the signed distance correlation and Pearson networks with the randomised networks occurs when we only use coexpression information. The *C*^†^ scores of *NS*(*d*_*S*_) and *NP* (*d*_*S*_) are 10 and 8.5 times higher than the expected ones for random networks. This is not surprising because we have built the networks using gene expression information. However, the coexpression data used to construct these networks is different to the data in STRING. The scores excluding the coexpression information *C*^‡^ are 3.7 and 3.2 times higher than in the random networks. This is remarkable because this assessment is performed on data that is completely different than the data used to construct the networks. Hence, by applying our pipeline to gene expression data, we can identify new types of relationships between proteins (and genes). These results demonstrate the power of gene coexpression networks to predict functional interactions, especially when constructed using signed distance correlation.

**Table 3:** Evaluation of the biological content of the networks with STRING. RE indicates the expected (mean) result based on random networks with the indicated edge density and its standard deviation.

| Network | All of STRING<br>information ($C$) | Only coexpression<br>information ($C^\dagger$) | All information<br>except coexpression ($C^\ddagger$) |
| --- | --- | --- | --- |
| $NS(d_S)$ | **9,277,087** | **2,825,233** | **8,318,736** |
| $NP(d_S)$ | 7,904,962 | 2,391,840 | 7,119,336 |
| RE $d_S$ | 2,348,373<br>$\pm 24,951$ | 281,405.5<br>$\pm 6,986$ | 2,223,120<br>$\pm 23,769$ |
| $NS(d_P)$ | **10,902,081** | **3,111,334** | **9,815,418** |
| $NP(d_P)$ | 9,490,978 | 2,690,505 | 8,578,437 |
| RE $d_P$ | 3,067,122<br>$\pm 33,510$ | 371,028.9<br>$\pm 10,515$ | 2,901,972<br>$\pm 31,648$ |

**Figure 4:**
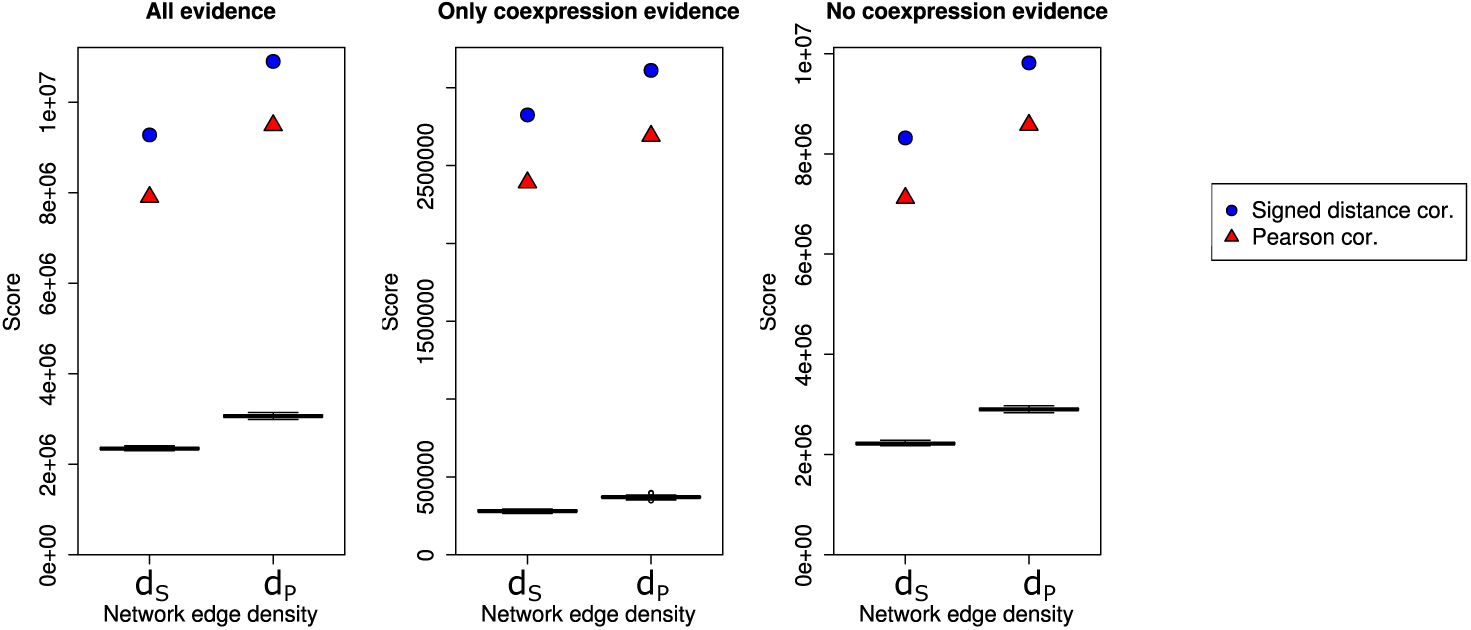
Scores obtained for the *R. leguminosarum* gene coexpression networks using STRING. All panels show the score for the different networks in the y-axis, and the network density on the x-axis. The scores are the result of adding up the confidence scores with all evidence (*C*), only coexpression evidence (*C*^†^) and everything excluding coexpression (*C*^‡^) from STRING associated with the edges in the networks, each computed using different of information. The black box plots correspond to the scores obtained by 30 random networks. Blue circles and red triangles represent signed distance correlation and Pearson correlation, respectively.

The analysis for networks with *d*_*P*_ tells a similar story; the scores obtained by the signed distance correlation network *NS*(*d*_*P*_) are higher than the scores from the Pearson correlation network *NP* (*d*_*P*_) despite *d*_*P*_ being the optimal edge density for Pearson correlation. We highlight the fact that when we use only coexpression information (*C*^†^), the score of *NS*(*d*_*S*_) is higher than the score of *NP* (*d*_*P*_), even though the former has fewer edges than the latter, and therefore its score is the result of adding fewer terms.

For the three datasets we analyse, signed distance correlation networks perform better than Pearson correlation networks with matching densities, according to both evaluations and for all tested edge densities.

## 4 Discussion

In this work we have introduced signed distance correlation, and presented a method to construct networks in a self-consistent way exclusively from gene expression data. This method has three main steps: data pre-processing, computing correlations, and thresholding. These steps combine well-established methods such as quantile normalisation, and the use of COGENT (Bozhilova *et al.*, 2020) to identify the optimal threshold.

Distance correlation is an intuitive approach to study gene expression because it relies on the differences in the expression between samples. By incorporating signs into distance correlation we can also differentiate between positive and negative relationships, and maintain the advantages of distance correlation.

We apply our method to data from *R. leguminosarum*, yeast, and human. In all cases our method produces networks that are more self-consistent than using Pearson correlation. The reason why self-consistency is so important is that it ensures that our results are robust to changes or noise in the data. Therefore, even when we cannot assess the biological significance of a network directly, we can be confident about it by measuring its self-consistency. Networks from signed distance correlation also capture more biological information than networks from Pearson correlation, as shown by our evaluation using STRING. In the case of *R. leguminosarum*, the signed distance correlation network (using an optimal threshold *θ*^*\**^ found with COGENT) captures almost four times more biological information than random networks. The amount of captured biological information is less if we use Pearson or Spearman correlations to build the networks (Fig. S1). These results suggest that self-consistent networks derived from biological data might capture more biological information than those with less stability. Therefore, even when we cannot assess the biological significance of a network directly we can be confident about it by measuring its self-consistency.

We applied our method to construct, to our knowledge, the first gene coexpression network for *R. leguminosarum*. This network promises to reveal rich biological information that will illuminate our understanding of plant-bacteria interactions and nitrogen fixation, and it is therefore the starting point for further investigations of the biological mechanisms of this organism.

Finally, we have showcased our method on gene expression datasets from different organisms obtained using different techniques: microarrays (*R. leguminosarum*), RNA-Seq (yeast), and single-cell RNA-Seq (human). However, the methods that we have developed are general, and can also be used to construct networks in a vast range of domains, such as, for example, economics (Wang *et al.*, 2018), neuroscience (Bernhardt *et al.*, 2011), climatology (Donges *et al.*, 2009), or any discipline where networks can be constructed from correlation data.

## Acknowledgements

We thank Alison K East and Florian Klimm for help and advice on how to analyse the datasets used in this manuscript. We thank Katharine Turner (ANU), Florian Klimm, and Malte D Luecken for fruitful discussions. In addition, we acknowledge support from COST Action CA15109, Keble College Oxford and Keble Association. The authors would like to acknowledge the use of the University of Oxford Advanced Research Computing (ARC) facility in carrying out this work (http://dx.doi.org/10.5281/zenodo.22558).

