## Supplementary Information for "Robust gene coexpression networks using signed distance correlation"

### 1 *Rhizobium leguminosarum* expression data

Table S1 provides the details of the gene expression data employed to generate the gene coexpression networks for *R. leguminosarum.* We used 54 different microarrays obtained under 18 conditions (first column). The second column indicates the growth condition of the bacteria; for example, "7 days pea, 1 day PI" stands for inoculation of seven days old pea plant sampled one day post-inoculation. The last column indicates the type of sample, which can be "Free-living" (bacteria grown in liquid culture), "Rhizosphere" (bacteria isolated from the fraction of soil in contact with the plant), or "Bacteroid" (bacteria that have infected the plant and differentiated). This data has been published previously [8, 6, 3] but never analysed jointly.

| Microarrays (channels) | Growth condition | Type |
|---|---|---|
| kk21 (Cy3), kk41 (Cy5), kk45 (Cy3) | Succinate $NH_4$ | Free-living |
| Jay10 (Cy5), Jay11 (Cy5), Jay12 (Cy3) | Glucose Glutamate | Free-living |
| Jay10 (Cy3), Jay11 (Cy3), Jay12 (Cy5) | Glucose Aspartate | Free-living |
| kk46 (Cy3), kk48 (Cy3), kk50 (Cy3) | Pyruvate $NH_4$ | Free-living |
| Vinoy43 (Cy5), Vinoy44 (Cy5), Vinoy45 (Cy3) | Pyruvate $NH_4$ Hespertin | Free-living |
| Ade7 (Cy3), Ade9 (Cy3), Ade18 (Cy5) | Pyruvate $NH_4$ IAA | Free-living |
| Ade8 (Cy5), Ade11 (Cy3), Ade15 (Cy5) | Pyruvate $NH_4$ Kinetin | Free-living |
| kk39 (Cy5), kk43 (Cy5), kk51 (Cy3) | Inositol $NH_4$ | Free-living |
| Vinoy26 (Cy5), Vinoy31 (Cy5), Vinoy34 (Cy3) | 7 days pea, 1 day PI | Rhizosphere |
| Vinoy27 (Cy5), Vinoy32 (Cy5), Vinoy35 (Cy3) | 7 days pea, 3 day PI | Rhizosphere |
| Vinoy28 (Cy5), Vinoy33 (Cy5), Vinoy36 (Cy3) | 7 days pea, 7 day PI | Rhizosphere |
| Vinoy29 (Cy5), Vinoy37 (Cy5), Vinoy39 (Cy3) | 14 days pea, 1 day PI | Rhizosphere |
| Vinoy30 (Cy5), Vinoy38 (Cy5), Vinoy40 (Cy3) | 21 days pea, 1 day PI | Rhizosphere |
| kk67 (Cy5), kk71 (Cy3), kk72 (Cy5) | 1 week pea, 7 days PI | Bacteroid |
| kk65 (Cy5), kk68 (Cy3), kk69 (Cy3) | 1 week pea, 15 days PI | Bacteroid |
| kk73 (Cy5), kk74 (Cy3), kk75 (Cy5) | 1 week pea, 21 days PI | Bacteroid |
| kk64 (Cy5), Vinoy22 (Cy3), Vinoy24 (Cy3) | 1 week pea, 28 days PI | Bacteroid |
| kk70 (Cy5), kk60 (Cy3), kk61 (Cy5) | Vetch seed, 28 days PI | Bacteroid |

Table 1: *Rhizobium leguminosarum* gene expression data description

#### 2 Spearman correlation networks for *R. leguminosarum*

We construct a Spearman correlation matrix $R$ from the pre-processed gene expression matrix $M^*$ by computing the Spearman correlation between every pair of gene expression vectors. We threshold $R$ to construct two networks $NR(d_S)$ and $NR(d_P)$ with edge density $d_S$ and $d_P$, respectively. Similarly to the networks obtained using Pearson correlation $NP(d_S)$ and $NP(d_P)$, the ones based on Spearman correlation have a larger and less densely connected largest connected component than $NS(d_S)$ and $NS(d_P)$. The summaries of these two networks can be found in Table S2. We evaluate the Spearman networks using STRING as described in the paper. The results obtained are higher than the obtained by the Pearson correlation networks but lower than those using signed distance correlation. Table S3 and Figure S1 show the results of the STRING evaluation for the Spearman networks.

| Network | Number of Edges | Number of vertices in LCC | Edge density LCC * 100 | Global clustering coeff. LCC |
|---|---|---|---|---|
| $NR(d_S)$ | 313,348 | 6,664 | 1.41 | 0.515 |
| $NR(d_P)$ | 406,977 | 6,831 | 1.34 | 0.515 |

Table 2: Summaries of Spearman networks for *R. leguminosarum*
LCC Denotes largest connected component.

| Network | All of STRING information ($C$) | Only coexpression information ($C^\dagger$) | All information except coexpression ($C^\ddagger$) |
|---|---|---|---|
| $NR(d_S)$ | 8,936,856 | 2,715,646 | 7,994,795 |
| $NR(d_P)$ | 10,488,856 | 3,003,020 | 9,422,007 |

Table 3: Evaluation of the biological content of the Spearman networks with STRING

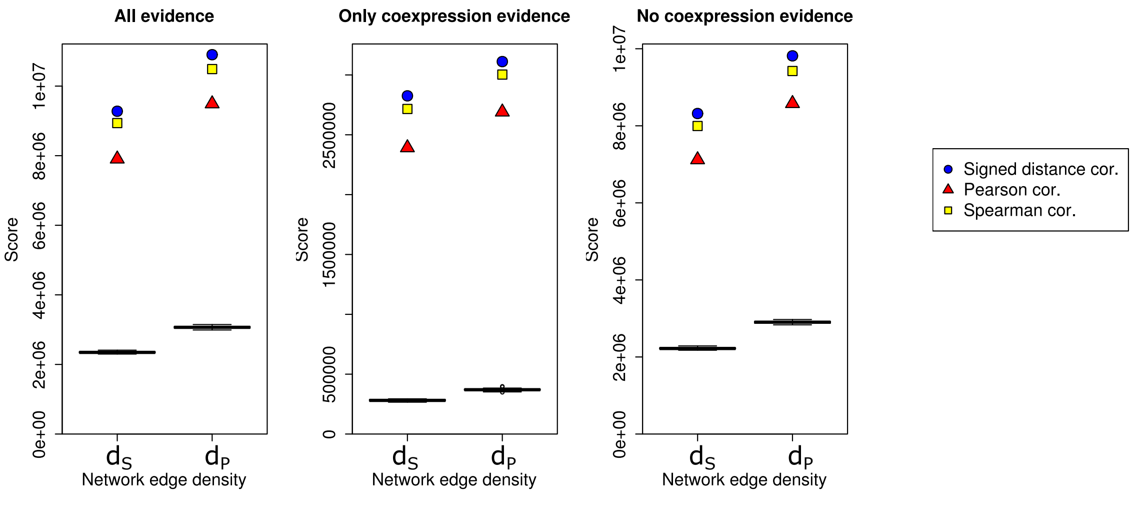


Figure 1: Scores obtained for the *R. leguminosarum* gene coexpression networks using STRING. All panels show the score for the different networks in the y-axis, and the network density on the x-axis. The scores are the result of adding up the confidence scores with all evidence ($C$), only coexpression evidence ($C^\dagger$) and everything excluding coexpression ($C^\ddagger$) from STRING associated with the edges in the networks, each computed using different of information. The black box plots correspond to the scores obtained by 30 random networks. Blue circles, red triangles and yellow squares represent signed distance correlation, Pearson correlation and Spearman correlation, respectively.

#### 3 Gene coexpression network analysis for the study of Yeast RNA-Seq data

We use our signed correlation pipeline to generate a gene coexpression network from a dataset obtained using RNA-Seq of yeast (*Saccharomyces cerevisiae*) expressing pathways designed to increase ATP or GTP consumption. We obtain all the raw-counts for experiment E-MTAB-5174 in Expression Atlas [5] and remove the genes with zero expression variance. The final dataset which we feed into our pipeline includes the expression of 6,930 genes across 209 samples.

Following the pipeline described in the methodology section, we pre-process the data and obtain the distance correlation matrix $D$, the Pearson correlation matrix $P$, the signed correlation matrix $S$, and the matrix $|P|$ of absolute values of the Pearson correlation. Table S4 and Figure S2 show the summaries and distribution of these matrices.

| Correlation matrix | Min | 1st q | Median | 3rd q | Max | Mean |
|---|---|---|---|---|---|---|
| $D$ | 0.00 | 0.26 | 0.40 | 0.57 | 1.00 | 0.42 |
| $\|P\|$ | 0.00 | 0.13 | 0.30 | 0.51 | 1.00 | 0.34 |
| $P$ | -0.96 | -0.29 | 0.00 | 0.31 | 1.00 | 0.01 |
| $S$ | -0.97 | -0.39 | -0.05 | 0.40 | 1.00 | 0.01 |

Table 4: Summaries for the correlation matrices of the Yeast dataset

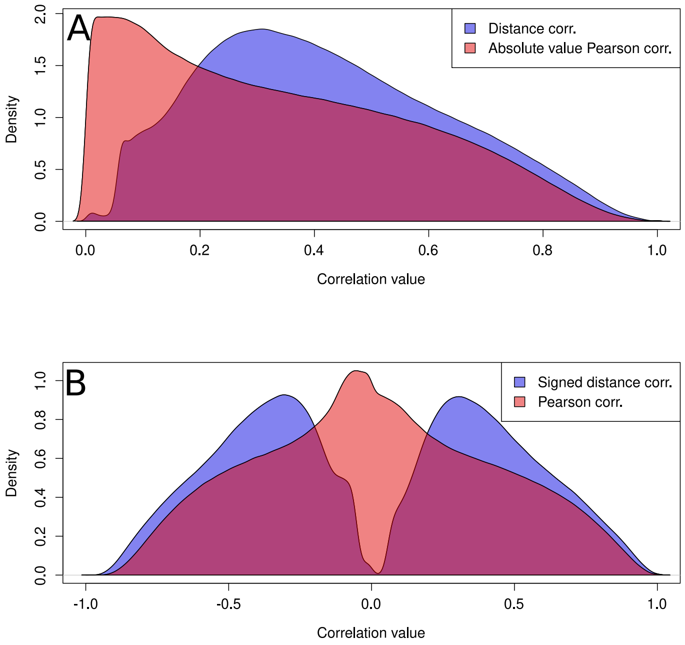


Figure 2: Density plots of the distribution of the values of the correlation matrices from the Yeast dataset. Panel A shows the distribution of values in the distance correlation matrix (blue) and of the absolute value of the values in the Pearson correlation matrix (red). Panel B shows the distribution of values in the signed distance correlation matrix (blue) and of the values in the Pearson correlation matrix (red).

We estimate the optimal threshold values $\theta^*$ and $\theta^\star$ to construct the unweighted gene coexpression networks $A_S(\theta^*)$ and $A_P(\theta^\star)$ using COGENT [2]. We follow our pipeline to calculate the self-consistency of the networks. We adjust the similarity score by subtracting the network density. Fig S3 shows the variation of the score function for the correlation matrices across different edge densities. For both signed distance correlation and Pearson correlation, there is a edge density value for which the score function reaches its maximum. This value is $d_S = 0.0131$ for the signed distance correlation (score of 0.746, which is achieved for $\theta^*$= 0.84) and $d_P = 0.0146$ for the Pearson correlation (score of 0.733, which is achieved for $\theta^\star = 0.81$). The results show that the use of signed distance correlation offers more stable networks. However, the difference between the performance of the two studied correlations is lower than the observed in the study of *R. leguminosarum*. We analyse the networks $NS(d_S)$ (edge density $d_S$) and $NS(d_P)$ (edge density $d_P$) retrieved from $S$, the networks $NP(d_S)$ (edge density $d_S$) and $NP(d_P)$ (edge density $d_P$) retrieved from $P$, and the networks $NR(d_S)$ (edge density $d_S$) and $NR(d_P)$ (edge density $d_P$) obtained using Spearman correlation. The summaries of the six networks are detailed in Table S5.

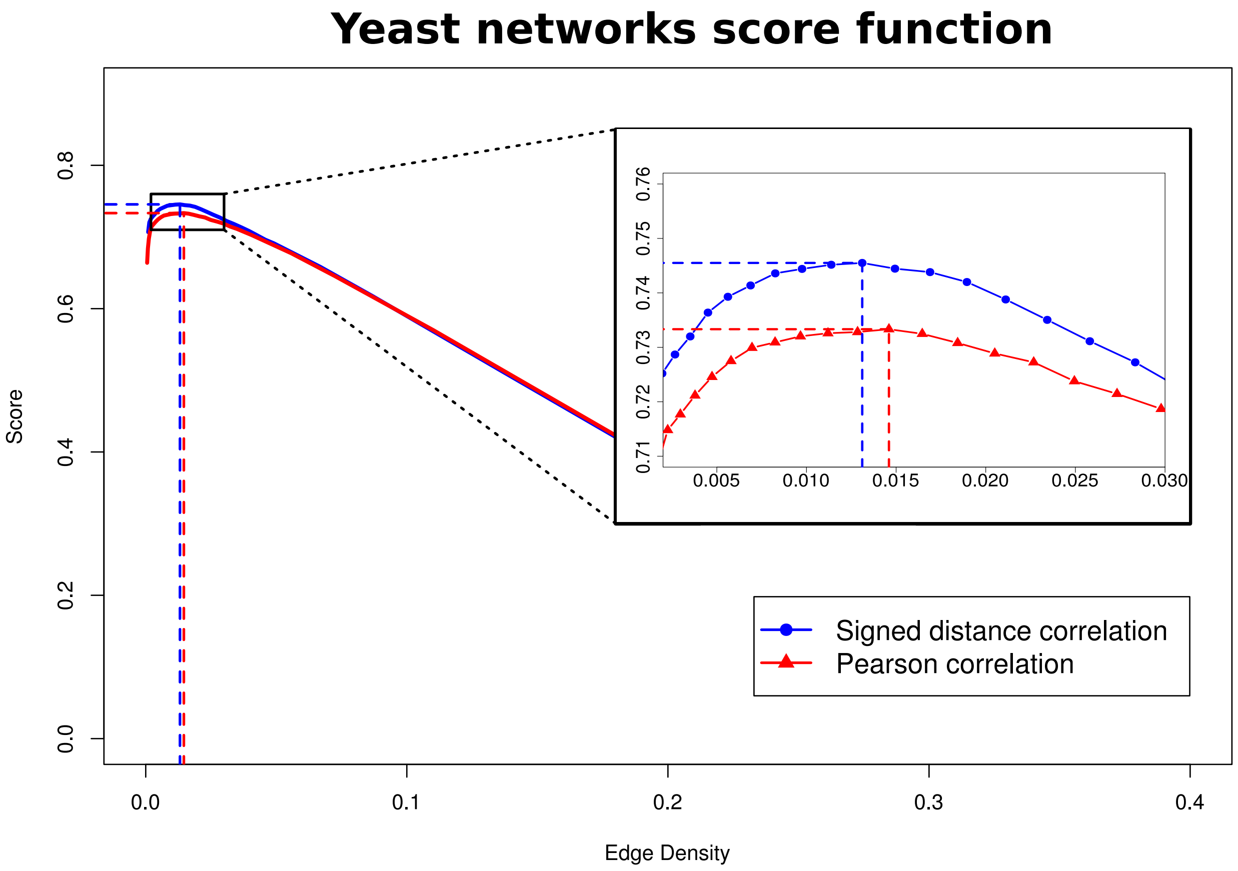


Figure 3: Score function values for different edge densities using the Yeast dataset. The blue line with circles shows the scores obtained using signed distance correlation and the red line with triangles those obtained using Pearson correlation. The dotted lines indicate the position of the highest score point for each line. This value is 0.746 for the signed distance correlation network (giving edge density 0.0131 which is achieved with $\theta^* = 0.84$) and 0.733 for the Pearson correlation network (giving edge density 0.0146 which is achieved with $\theta^\star = 0.81$).

We evaluate the six networks using STRING [9]. We use the three different sets of confidence scores: obtained using all the evidence in STRING ($C$), obtained using only coexpression information ($C^\dagger$), and obtained using all the evidence except the coexpression information ($C^\ddagger$). Table S6 and Figure S4 present the results. For all the studied cases, the results obtained using the signed distance correlation networks are the highest. Unlike with the *R. leguminosarum* dataset, the Pearson correlation networks perform better than the Spearman correlation networks. In fact, the results for the signed distance correlation and Pearson correlation are almost identical (as in the case of the self-consistency study).

| Network | Number of Edges | Number of vertices in LCC | Edge density LCC * 100 | Global clustering coeff. LCC |
|---|---|---|---|---|
| $NS(d_S)$ | 314,771 | 3,987 | 3.91 | 0.651 |
| $NS(d_P)$ | 350,549 | 4,154 | 4.02 | 0.607 |
| $NP(d_S)$ | 314,771 | 4,497 | 3.07 | 0.431 |
| $NP(d_P)$ | 350,549 | 4,615 | 3.25 | 0.516 |
| $NR(d_S)$ | 314,771 | 5,060 | 2.39 | 0.578 |
| $NR(d_P)$ | 350,549 | 5,117 | 2.44 | 0.578 |

Table 5: Summaries of yeast networks. LCC denotes largest connected component.

These results suggest that a high similarity in the self-consistency of the networks may imply a similarity in the amount of biological information that the networks are able to capture.

We generate 60 random networks of which 30 have edge density $d_S$ and the other 30 have edge density $d_P$. We evaluate these networks with the STRING information and find that the six networks in Table S5 outperform all of them. The highest difference to random is obtained for signed distance correlation networks when using only coexpression information ($C^{\dagger}$) to evaluate the networks; for $NS(d_S)$ the score is 17.69 times higher than the mean score obtained by random networks with matching densities.

| Network | All of STRING information ($C$) | Only coexpression information ($C^{\dagger}$) | All information except coexpression ($C^{\ddagger}$) |
|---|---|---|---|
| $NS(d_S)$ | **36,941,775** | **26,763,561** | **21,135,268** |
| $NP(d_S)$ | 36,610,618 | 26,706,960 | 20,873,493 |
| $NR(d_S)$ | 23,557,830 | 15,143,418 | 15,889,580 |
| RE $d_S$ | 4,412,687 ± 43,162 | 1,512,629 ± 24,840 | 3,741,742 ± 36,649 |
| $NS(d_P)$ | **39,013,837** | **27,935,541** | **22,642,609** |
| $NP(d_P)$ | 38,609,494 | 27,822,217 | 22,310,410 |
| $NR(d_P)$ | 25,395,572 | 16,143,245 | 17,243,212 |
| RE $d_P$ | 4,916,218 ± 50,112 | 1,678,870 ± 24,502 | 4,174,943 ± 46,196 |

Table 6: Evaluation of the biological content of the networks with STRING. RE indicates the expected (mean) result based on random networks with the indicated edge density and its standard deviation.

We conclude that the Yeast gene coexpression networks obtained using signed distance correlation are more stable and recover more biological information than those based on Pearson correlation or Spearman correlation.

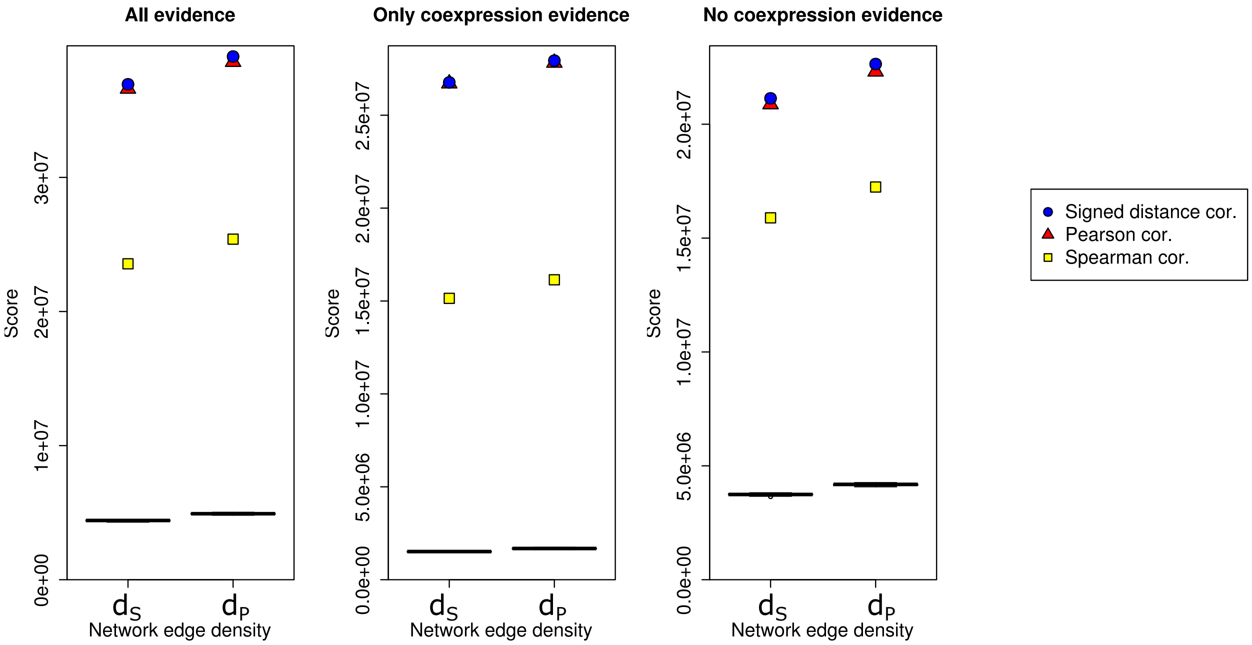


Figure 4: Scores obtained for the yeast gene coexpression networks using STRING. All panels show the score for the different networks in the y-axis, and the network density on the x-axis. The scores are the result of adding up the confidence scores from STRING associated with the edges in the networks. Each plot corresponds to a different set of values: using all evidence $C$, only coexpression evidence $C^{\dagger}$ and excluding coexpression evidence $C^{\ddagger}$. The black box plots correspond to the scores obtained by 30 random networks. Blue circles, red triangles and yellow squares represent signed distance correlation, Pearson correlation and Spearman correlation, respectively.

#### 4 Gene coexpression network analysis for the study of human liver single-cell RNA-Seq data

We use our signed correlation pipeline to generate a gene coexpression network from a dataset obtained using single-cell RNA-Seq of human liver cells [4]. The original dataset measures the expression of 15,353 genes in 1,622 cells.

This dataset is considerably different from the previous two, both in the data itself – the measurements correspond to different cells instead of to different samples – and the organism – while both *R. leguminosarum* and *S. cerevisiae* are unicellular organisms, humans are not. Hence, a considerable proportion of genes are not expressed in the studied cells (for example, specific genes in neurons will not be expressed in cells from liver). For this reason, we employ a different pre-processing strategy in this case. In the first place, we quantile-normalise the data [1] to make the measurements in the different cells comparable. Afterwards, as in [7], we identify the "non-changing genes". These genes are those for which the difference between its highest and lowest expression value ("expression difference") is lower than the median of all the expression differences calculated for each gene, and in addition for which the mean expression signal between samples is lower than the median of all the expression signals calculated for each gene. After removing the "non-changing genes" we obtain an already quantile-normalised dataset with information for 8,585 genes. We do not apply more preprocessing steps to this dataset (we do not quantile normalise again the data and we do not set the lowest expressed genes from each sample to the lowest expression value).

We calculate the distance correlation matrix ($D$), the Pearson correlation matrix ($P$), the signed correlation matrix ($S$), and the matrix $|P|$ of absolute values of the Pearson correlation. Table S7 and Figure S5 show the summaries and distribution of these matrices.

| Correlation matrix | Min | 1st q | Median | 3rd q | Max | Mean |
|---|---|---|---|---|---|---|
| $D$ | 0.00 | 0.03 | 0.04 | 0.05 | 0.88 | 0.04 |
| $\|P\|$ | 0.00 | 0.01 | 0.02 | 0.03 | 0.91 | 0.02 |
| $P$ | -0.63 | -0.02 | 0.00 | 0.02 | 0.91 | 0.00 |
| $S$ | -0.71 | -0.03 | -0.02 | 0.04 | 0.88 | 0.00 |

Table 7: Summaries for the correlation matrices of the single-cell dataset from human liver

We estimate the optimal threshold values $\theta^*$ and $\theta^\star$ to construct unweighted gene coexpression networks $A_S(\theta^*)$ and $A_P(\theta^\star)$ using COGENT [2]. We follow a similar pipeline as the one employed in the case of *R. leguminosarum* to calculate and adjust the self-consistency of the networks. The only difference is that as the input expression matrix is already quantile-normalised and low expressed genes have already been filtered out, in each of the 25 iterations the pre-processing steps are omitted. Figure S6 shows the variation of the score function for the correlation matrices across different edge densities. In both cases, there is a edge density value for which the score function reaches its maximum. This value is $d_S = 0.00009$ for the signed distance correlation (score of 0.896, which is achieved for $\theta^* = 0.44$) and $d_P = 0.000089$ for the Pearson correlation (score of 0.781, which is achieved for $\theta^\star = 0.43$). The results show that for all tested edge densities, the use of signed distance correlation offers more stable networks. We analyse the networks $NS(d_P)$ (edge density $d_S$) and $NS(d_P)$ (edge density $d_P$) retrieved from $S$, the networks $NP(d_S)$ (edge density $d_S$) and $NP(d_P)$ (edge density $d_P$) retrieved from $P$. We also construct two networks from a correlation matrix obtained using Spearman correlation: $NR(d_S)$ (edge density $d_S$) and $NR(d_P)$ (edge density $d_P$). The summaries of the six networks are detailed in Table S8. We note that the largest connected components of the networks contain a lower proportion of the total vertices than in previous datasets. The networks obtained using signed distance correlation have a smaller and denser largest connected component than those obtained using Pearson correlation. In contrast to the two previous datasets, the largest connected component of the networks obtained using Spearman correlation is smaller and denser than those from signed distance correlation and Pearson correlation networks; however, the signed distance correlation networks still have the highest global clustering coefficient.

Similarly as for the two previous datasets, we evaluate the constructed networks using three sets of

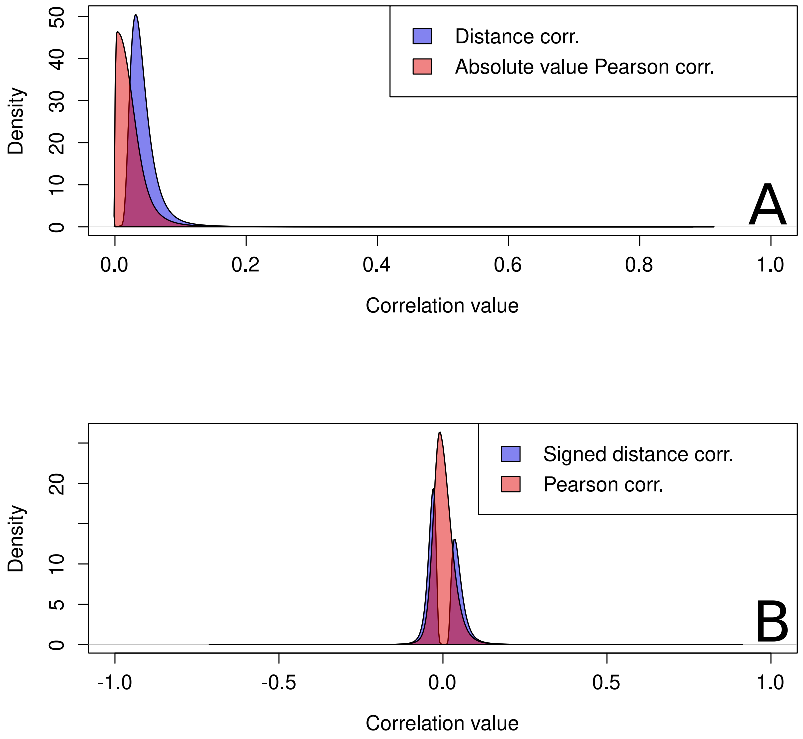


Figure 5: Density plots of the distribution of the values of the correlation matrices from the Yeast dataset. Panel A shows the distribution of values in the distance correlation matrix (blue) and of the absolute value of the values in the Pearson correlation matrix (red). Panel B shows the distribution of values in the signed distance correlation matrix (blue) and of the values in the Pearson correlation matrix (red).

| Network | Number of Edges | Number of vertices in LCC | Edge density LCC * 100 | Global clustering coeff. LCC |
|---|---|---|---|---|
| $NS(d_S)$ | 3320 | 212 | 14.61 | 0.87 |
| $NS(d_P)$ | 3279 | 210 | 14.71 | 0.87 |
| $NP(d_S)$ | 3320 | 219 | 12.42 | 0.83 |
| $NP(d_P)$ | 3279 | 216 | 12.62 | 0.83 |
| $NR(d_S)$ | 3320 | 161 | 25.54 | 0.81 |
| $NR(d_P)$ | 3279 | 160 | 25.54 | 0.81 |

Table 8: Summaries of single cell human liver networks.
LCC denotes largest connected component.

confidence values obtained from STRING. We also evaluate 60 random networks with edge densities $d_S$ (30 networks) and $d_P$ (30 networks). Table S9 and Figure S7 show the results. The networks obtained using Spearman correlation get a higher score than the Pearson ones when evaluating them using exclusively the coexpression information from STRING ($C^\dagger$), whereas the situation is the reverse in the two other scenarios ($C$ and $C^\ddagger$). Overall, the networks obtained using signed distance correlation $NS(d_S)$ and $NS(d_P)$ retrieve a higher score than their competitors based on either Pearson or Spearman correlation. This result suggest that among the assessed methods, signed distance correlation captures the broadest range of biological information, which makes it a tool of choice to generating gene coexpression networks. All the analysed networks obtain a higher score than the random ones. The highest difference to random is obtained for signed distance correlation networks when using only coexpression information ($C^\dagger$) to evaluate the networks; for $NS(d_P)$ the score is 92.92 times higher than the mean score obtained by random networks with matching densities; for $NS(d_S)$ it is even 94.28 than the corresponding mean score for the random networks.

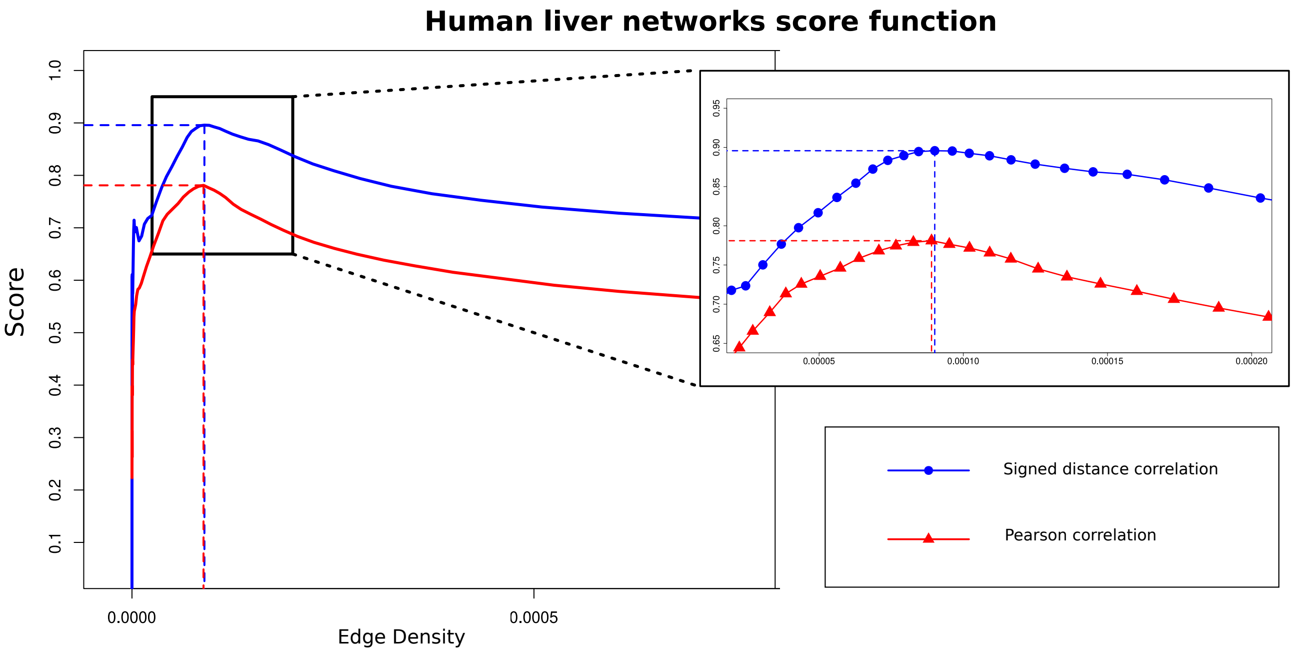


Figure 6: Plot of the score function values for different edge densities using the human liver single-cell RNA-Seq dataset. The blue line shows the scores obtained using signed distance correlation and the red line those obtained using Pearson correlation. The dotted lines indicate the position of the highest score point for each line. This value is 0.896 for the signed distance correlation network (giving edge density $9.01e^{-05}$ which is achieved with $\theta^* = 0.44$) and 0.781 for the Pearson correlation network (giving edge density $8.90e^{-05}$ which is achieved with $\theta^* = 0.43$).

| Network | All of STRING information ($C$) | Only coexpression information ($C^\dagger$) | All information except coexpression ($C^\ddagger$) |
|---|---|---|---|
| $NS(d_S)$ | **703,908** | **615,363** | **675,788** |
| $NP(d_S)$ | 651,712 | 563,881 | 624,554 |
| $NR(d_S)$ | 607,793 | 574,426 | 595,682 |
| RE $d_S$ | 22,398 ± 2,434 | 6,527 ± 971 | 19,593 ± 2,430 |
| $NS(d_P)$ | **696,299** | **609,262** | **668,927** |
| $NP(d_P)$ | 645,745 | 560,246 | 619,247 |
| $NR(d_P)$ | 603,929 | 570,977 | 592,232 |
| RE $d_P$ | 232,378 ± 2,687 | 6,557 ± 1,064 | 19,605 ± 2,552 |

Table 9: Evaluation of the biological content of the networks with STRING. RE indicates the expected (mean) result based on random networks with
the indicated edge density and its standard deviation.

We conclude that the liver gene coexpression networks obtained using signed distance correlation are more stable and show recover more biological information than those based on Pearson correlation and that our signed distance correlation method is suitable for studying single cell gene expression data.

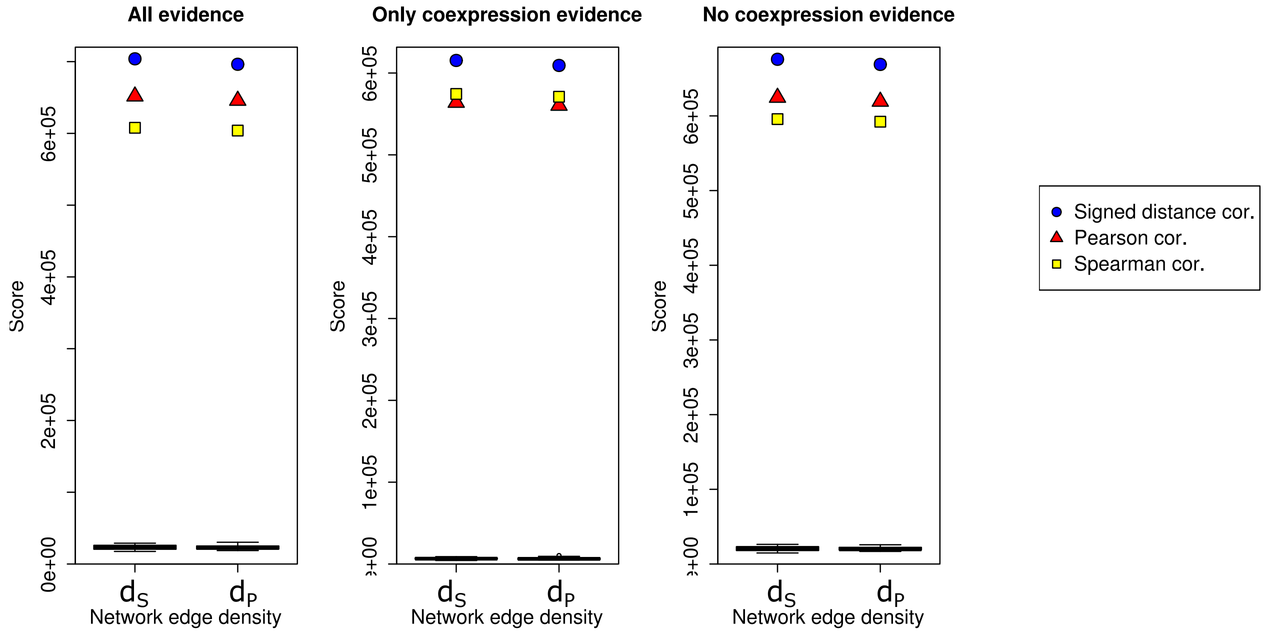


Figure 7: Scores obtained for the single-cell human liver gene coexpression networks using STRING. All panels show the score for the different networks in the y-axis, and the network density on the x-axis. The scores are the result of adding up the confidence scores from STRING associated with the edges in the networks. Each plot corresponds to a different set of values: using all evidence $C$, only coexpression evidence $C^{\dagger}$ and excluding coexpression evidence $C^{\ddagger}$. The black box plots correspond to the scores obtained by 30 random networks. Blue circles, red triangles and yellow squares represent signed distance correlation, Pearson correlation and Spearman correlation, respectively.

#### 5 Network evaluation using information from STRING

We use the interactions between proteins reported in STRING to evaluate the amount of biological information that the constructed networks capture. The association evidence in STRING is categorized into independent channels, weighted, and integrated, resulting in a confidence score for all recorded protein interactions. For each organism, we use three different sets of interactions and confidence scores: the overall confidence score $C$ provided by STRING, the $C^{\dagger}$ obtained attending only to coexpression information, and the $C^{\ddagger}$ obtained excluding all coexpression information.

The recomputing of the each of the scores was done using the python script located on the STRING webpage (Figure 8), which was accessed on the 20$^{\text{th}}$ of April 2020 using the URL https://string-db.org/cgi/help.pl?&subpage=faq%23how-are-the-scores-computed. The script can be found at the end of this section. We commented out lines 96–98, 100–102 in the script to compute the $C^{\dagger}$ values; and the line 99 in the script to obtain the $C^{\ddagger}$ values.

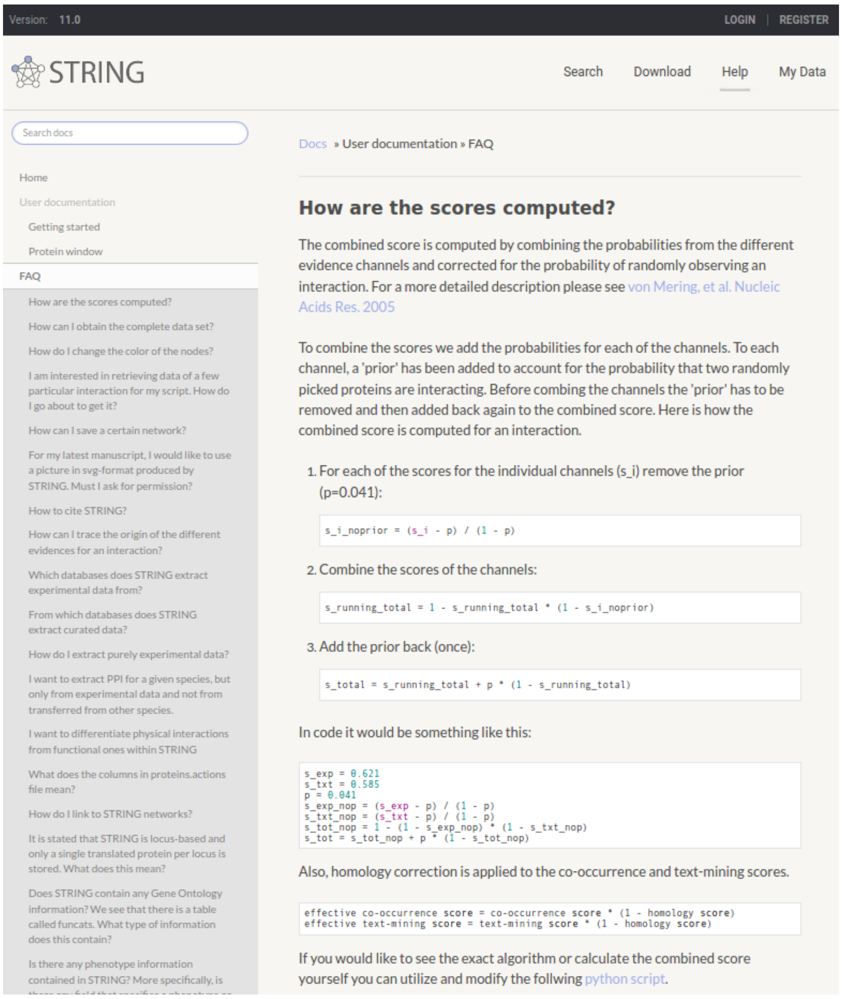


Figure 8: STRING FAQ webpage. Accessed on the 20$^{\text{th}}$ of April 2020. The content describes how the STRING combines the scores of the different channels. At the end of the section there is the link to the script which was modified and used in this work.

```python
from __future__ import print_function
import os
import sys

###########################################################
## This script combines all the STRING's channels subscores
## into the final combined STRING score.
## It uses unpacked protein.links.full.xx.txt.gz as input
## which can be downloaded from the download subpage:
##      https://string-db.org/cgi/download.pl
###########################################################

input_file = "9606.protein.links.full.v10.5.txt"

if not os.path.exists(input_file):
    sys.exit("Can't locate input file %s" % input_file)

prior = 0.041

def compute_prior_away(score, prior):

    if score < prior: score = prior
    score_no_prior = (score - prior) / (1 - prior)

    return score_no_prior

header = True
for line in open(input_file):

    if header:
        header = False
        continue

    l = line.split()

    ## load the line

    (protein1, protein2,
     neighborhood, neighborhood_transferred,
     fusion, cooccurrence,
     homology,
     coexpression, coexpression_transferred,
     experiments, experiments_transferred,
     database, database_transferred,
     textmining, textmining_transferred,
     initial_combined) = l

    ## divide by 1000

    neighborhood = float(neighborhood) / 1000
    neighborhood_transferred = float(neighborhood_transferred) / 1000
    fusion = float(fusion) / 1000
    cooccurrence =  float(cooccurrence) / 1000
    homology = float(homology) / 1000
    coexpression = float(coexpression) / 1000
    coexpression_transferred = float(coexpression_transferred) / 1000
    experiments = float(experiments) / 1000
    experiments_transferred = float(experiments_transferred) / 1000
    database = float(database) / 1000
    database_transferred = float(database_transferred) / 1000
    textmining = float(textmining) / 1000
    textmining_transferred = float(textmining_transferred) / 1000
    initial_combined = int(initial_combined)

    ## compute prior away

```

```python
    neighborhood_prior_corrected              = compute_prior_away (neighborhood, prior)
    neighborhood_transferred_prior_corrected = compute_prior_away (neighborhood_transferred, prior)
    fusion_prior_corrected                    = compute_prior_away (fusion, prior)
    cooccurrence_prior_corrected              = compute_prior_away (cooccurrence, prior)
    coexpression_prior_corrected              = compute_prior_away (coexpression, prior)
    coexpression_transferred_prior_corrected = compute_prior_away (coexpression_transferred, prior)
    experiments_prior_corrected               = compute_prior_away (experiments, prior)
    experiments_transferred_prior_corrected  = compute_prior_away (experiments_transferred, prior)
    database_prior_corrected                  = compute_prior_away (database, prior)
    database_transferred_prior_corrected     = compute_prior_away (database_transferred, prior)
    textmining_prior_corrected                = compute_prior_away (textmining, prior)
    textmining_transferred_prior_corrected   = compute_prior_away (textmining_transferred, prior)

    ## then, combine the direct and transferred scores for each category:

    neighborhood_both_prior_corrected = 1.0 - (1.0 - neighborhood_prior_corrected) * (1.0 - neighborhood_transferred_prior_corrected)
    coexpression_both_prior_corrected = 1.0 - (1.0 - coexpression_prior_corrected) * (1.0 - coexpression_transferred_prior_corrected)
    experiments_both_prior_corrected  = 1.0 - (1.0 - experiments_prior_corrected) * (1.0 - experiments_transferred_prior_corrected)
    database_both_prior_corrected     = 1.0 - (1.0 - database_prior_corrected) * (1.0 - database_transferred_prior_corrected)
    textmining_both_prior_corrected   = 1.0 - (1.0 - textmining_prior_corrected) * (1.0 - textmining_transferred_prior_corrected)

    ## now, do the homology correction on cooccurrence and textmining:

    cooccurrence_prior_homology_corrected = cooccurrence_prior_corrected * (1.0 - homology)
    textmining_both_prior_homology_corrected = textmining_both_prior_corrected * (1.0 - homology)

    ## next, do the 1 - multiplication:

    combined_score_one_minus = (
        (1.0 - neighborhood_both_prior_corrected) *
        (1.0 - fusion_prior_corrected) *
        (1.0 - cooccurrence_prior_homology_corrected) *
        (1.0 - coexpression_both_prior_corrected) *
        (1.0 - experiments_both_prior_corrected) *
        (1.0 - database_both_prior_corrected) *
        (1.0 - textmining_both_prior_homology_corrected) *
        1)

    ## and lastly, do the 1 - conversion again, and put back the prior *exactly once*

    combined_score = (1.0 - combined_score_one_minus)            ## 1- conversion
    combined_score *= (1.0 - prior)                               ## scale down
    combined_score += prior                                       ## and add prior.

    ## round

    combined_score = int(combined_score * 1000)
    print(protein1, protein2, combined_score)
```

Listing 1: Combine subscores script

[4] Florian Klimm, Enrique M Toledo, Thomas Monfeuga, Fang Zhang, Charlotte M Deane, and Gesine Reinert. Functional module detection through integration of single-cell rna sequencing data with protein–protein interaction networks. *bioRxiv*, page 698647, 2019.
